# Corticostriatal Beta Power Changes Associated with Cognitive Function in Parkinson’s Disease

**DOI:** 10.1101/2022.07.07.499165

**Authors:** DL Paulo, H Qian, D Subramanian, GW Johnson, K Hett, C Kao, N Roy, K Dhima, DO Claassen, SK Bick

**Affiliations:** Vanderbilt University Medical Center

## Abstract

Cognitive impairment (CI) is the most frequent nonmotor symptom in Parkinson’s Disease (PD) and is associated with deficits in executive functions such as working memory. Previous studies have demonstrated that caudate beta power is involved in learning and working memory. Decreased dopamine in motor cortico-striato-thalamo-cortical (CSTC) circuits results in increased beta power and PD motor symptoms. Analogous changes in cognitive CSTC circuits, including the caudate and dorsolateral prefrontal cortex (DLPFC), may contribute to PD CI. The objective of our study is to evaluate whether beta power changes in caudate and DLPFC contribute to cognitive impairment in PD patients. To investigate this, we used local field potential (LFP) recordings during deep brain stimulation surgery in 15 PD patients. LFP signals from DLPFC and caudate were performed at rest and during a verbal working memory task. We examined beta power changes during the working memory task and relationship of beta power to pre-operative neuropsychological testing results. Beta power decreased in both DLPFC and caudate during encoding of correct trials, whereas beta power increased in DLPFC and caudate during feedback for correct responses. Subjects with cognitive impairment showed smaller decreases in caudate and DLPFC beta power during encoding, greater increases in beta power during feedback, and lower average resting-state beta power. Additionally, reduced caudate beta power during encoding correlated with better memory scores on pre-operative neuropsychological testing, while greater DLPFC beta power during feedback correlated with worse scores in the attention domain. Our findings suggest that similar to the relationship between beta power in motor CSTC circuits and PD motor symptoms, beta power changes in parallel cognitive CSTC circuits may be correlated with cognitive symptoms in PD patients.

## Introduction

Parkinson’s disease (PD) is a movement disorder that affects approximately 1 million people in the US and 10 million people worldwide, and is expected to double in prevalence by 2030.^1^ Although PD is primarily thought of as a movement disorder with cardinal symptoms of bradykinesia, rigidity and tremor, nonmotor symptoms are also very common and have a significant impact on quality of life. Cognitive impairment (CI) is the most common nonmotor symptom, present in approximately 9-19% of all PD patients within 2 years of initial diagnosis^2–4^, 50% at 5-10 years after diagnosis ^5,6^, and up to 75% of patients at the time of death. PD CI is a major contributor to patient disability, decreased quality of life, and is associated with increased disease-related mortality.^7^

PD CI can involve deficits in heterogeneous cognitive domains including memory, attention, executive function, visuospatial function, and language. Executive function (including working memory), attention, and memory are most prominently affected in early PD,^8–10^ while longitudinal studies of PD CI have shown that significant declines in visuospatial function, memory, and global cognitive ability occur by mid-stage of the disease.^11–13^ Current first-line therapies for PD CI include behavioral therapy, acetylcholinesterase inhibitors, and memantine, however existing treatments unfortunately have limited efficacy. Improved understanding of the pathophysiology underlying PD CI is needed to support development of novel treatment modalities.

Imaging studies have demonstrated that the caudate is involved in cognitive changes in PD. Structural MRI and PET studies have demonstrated that caudate volume and dopamine levels inversely correlate with the degree of PD CI.^14,15^ Studies using functional MRI have demonstrated decreased caudate activity and decreased caudate-dorsolateral prefrontal cortex (DLPFC) connectivity in PD patients with CI.^16,17^ PD patients with CI have also been shown to have decreased caudate and DLPFC activity during a working memory task,^16^ with caudate activity specifically being decreased during working memory encoding.^18^

Observed involvement of caudate and DLPFC in PD CI may be related to dopamine-modulated changes in cortico-striato-thalamo-cortical (CSTC) circuits in PD. The striatum, which includes the nucleus accumbens, caudate, and putamen, is somatotopically organized into parallel CSTC circuits which contribute to motor, cognitive and limbic functions.^19^ PD motor symptoms have been associated with beta range (15-30 Hz) neural signal oscillation changes in motor CSTC structures including subthalamic nucleus, globus pallidus internus, and motor cortex.^20,21^ These beta changes are caused by decreased dopaminergic input to putamen^22^ and are reversed by medical and surgical therapies that improve PD motor symptoms.^23–27^ The cognitive CSTC circuit includes caudate and DLPFC and similarly has decreased dopaminergic input in PD. Beta power in caudate and DLPFC has also been associated with cognitive functions. Primate studies have shown that DLPFC beta bursting and power are suppressed during working memory encoding, while gamma (30-100 Hz) power and bursts are induced.^28–30^ Previous research in human subjects participating in a learning task found increased caudate and DLPFC beta power during feedback following correct trials, with increased DLPFC beta power correlating with learning.^31^ Whether beta power changes in cognitive CSTC circuits contribute to PD CI remains unknown.

The objective of this study was to determine whether caudate and DLPFC beta power changes contribute to cognitive impairment in PD. We performed local field potential (LFP) recordings from caudate and DLPFC in PD patients during deep brain stimulation (DBS) surgery at rest and during a working memory task. We report beta power changes during working memory that differ in patients with CI and correlate with preoperative memory domain function. These findings increase our understanding of the pathophysiology of PD CI and may provide the basis for future development of novel neuromodulation strategies.

## Methods

### Subjects

Fifteen PD patients undergoing DBS surgery of the subthalamic nucleus (STN) or globus pallidus internus (GPi) under local anesthesia participated in this study. All PD patients undergoing new DBS implantation, regardless of baseline cognitive status, were eligible to participate. Subjects provided written informed consent prior to participation. This prospective study was approved by the Vanderbilt University Medical Center Institutional Review Board prior to initiation. Demographic, behavioral, and clinical data from all subjects were collected from the electronic medical record.

### Preoperative Evaluation

As part of the routine DBS preoperative workup at our center all subjects underwent preoperative MRI brain scans and underwent formal motor and neuropsychological testing. Motor function on and off PD medications was graded by the Unified Parkinson’s Disease Rating Scale (UPDRS). Neuropsychological evaluation included tests to measure functioning within cognitive domains frequently impaired in PD: executive functioning, processing speed, attention, memory, visuospatial, and language. These tests included assessments of premorbid functioning, the Montreal Cognitive Assessment (MoCA), the Wechsler Adult Intelligence Scale (WAIS) IV Digit Span Test, Trail Making Tests A and B, the Delis–Kaplan Executive Function System (D-KEFS) Color-Word Interference Test, the Action Fluency Test, the F-A-S Letter Fluency Test, the Animals Category Fluency Test, the Neuropsychological Assessment Battery (NAB) Naming Test (Form 1), the NAB Visual Discrimination Test (Form 1), the Repeatable Battery for the Assessment of Neuropsychological Status (RBANS) Line Orientation Test (Form A), the Rey Auditory Verbal Learning Test, and the Wechsler Memory Scale (Fourth Edition Logical Memory).

For subjects who consented to participate in this research study, the planned DBS electrode trajectory was examined in the clinical planning software (Waypoint, FHC), and the relationship of the trajectory to caudate and DLPFC was noted. For trajectories which contacted caudate, the distance of this contact above the planned target was noted in order to determine the depth at which to perform intraoperative research recordings.

### Surgery

All patients who participated in this study underwent DBS electrode implantation surgery under local anesthesia with microelectrode recordings to help refine final electrode placement. At our center we generally use 3 microelectrode recording tracts per side implanted. Dopaminergic medications were held the night prior to surgery according to standard clinical protocol to facilitate intraoperative motor testing. A custom-made mini-stereotactic frame (FHC Inc., Bowdoin, ME) was mounted with two microdrives and microelectrodes were advanced to the structure to be recorded from (DLPFC or caudate) along the planned clinical trajectory to the treatment target. This setup allowed for simultaneous bilateral recordings. LFP recordings were performed from the macrocontact of the clinical microelectrodes and recorded via the FHC Guideline 5 system with a sampling rate of 1000Hz. Nine subjects had recording electrodes that traversed the caudate, and thirteen subjects had electrodes that traversed the DLPFC.

### Task

At the beginning of the research session, two minutes of LFP data was recorded while subjects rested quietly with their eyes closed.^32^ Subjects subsequently participated in a verbal 2-back working memory task ^33^, during which they were sequentially visually presented with a series of words and prompted to respond whether the word presented during the current trial matched the word presented two trials prior by pressing a button. Following a response, they were given visual feedback on whether their response was correct or incorrect **(Figure 1)**.

**Figure 1.**
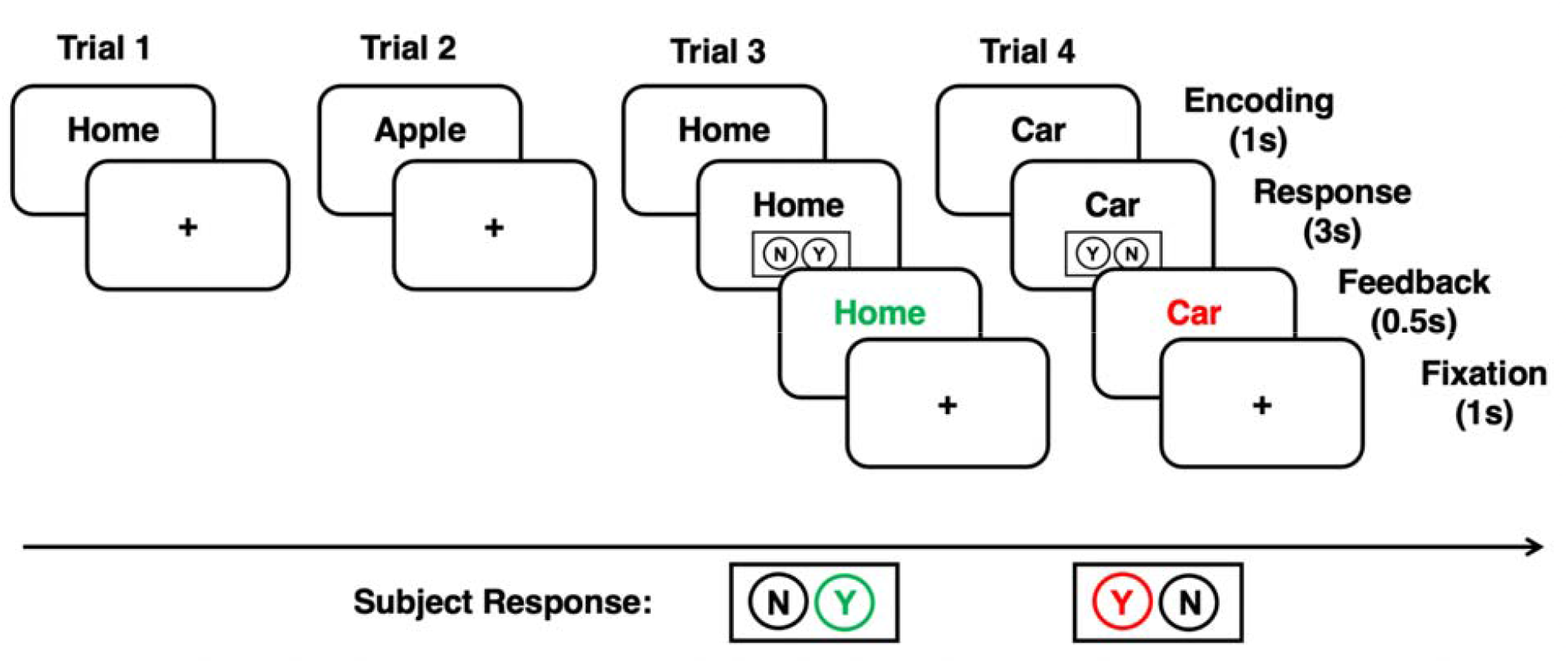
Depiction of 2-back Verbal Working Memory Task. The subject is presented with a word, and after a pause, a response cue will appear prompting the subject to answer whether the current word matches that from 2 trials perior. Following the response, visual feedback will be presented. Green represents a correct response, red represents an incorrect response. Y=yes, N=no.

Subjects completed two 75-trial blocks of this task. The task was run using MonkeyLogic task presentation software (NIMH MonkeyLogic), with transistor-transistor logic (TTL) pulses sent to the FHC Guideline 5 neurophysiology system to allow alignment of task events with neural recordings.

### Neuropsychological testing data

Preoperative neuropsychological testing measures were categorized into the cognitive domains of executive function, attention, processing speed, memory, language, or visuospatial function (**Supplementary Table 1**). For each subject, we converted their norm-referenced standardized assessment scores to Z-scores and averaged within each cognitive domain to create a domain-specific composite score.

We categorized patients as cognitively impaired if they had impairment on at least two individual task measures, per the Movement Disorder Society (MDS) Task Force’s level II criteria for PD mild cognitive impairment (MCI).^34^ We used the Wechsler cutoff (<9th percentile) to classify impairment on neuropsychological test measures.

### Imaging

Following surgery, patients underwent postoperative head CT as part of standard clinical protocol. This was merged with their preoperative MRI using Waypoint software to evaluate final electrode placement and confirm relative positioning of recording microelectrodes. Any microelectrode tracts that were determined by two neurosurgeons (DP, SB) to be out of the region of interest for this study were excluded from analysis.

Given the previously reported relationship between caudate volume and cognition in PD patients, we computed caudate volume to use as a covariate in our analyses. Estimation of caudate volumes was obtained by feeding subjects’ T_1_-weighted MRI to a recent deep-learning approach based on a large ensemble of fully convolutional neural networks.^35^ This method provides automatic parcellation of 133 brain structures following the Desikan-Killiany-Tourville protocol.^36^ All caudate volumes were given in the T_1_-weighted native space after total intracranial volume normalization.

### Neurophysiology and Statistical Analysis

LFP analysis was performed offline using MATLAB (MathWorks, Natick MA) and Fieldtrip MATLAB toolbox.^37^ Channels were visually examined for noise and excluded if the signal was contaminated by significant artifact. Data was bandpass filtered at 60 Hz to remove line noise, demeaned by channel, and aligned to task events using digital event triggers. Beta power was calculated using Morlet wavelet time-frequency transformation in Fieldtrip, and task-based beta power was Z-scored for each subject, channel, and frequency across all trials.

To examine working memory related beta power changes, we divided the task into several epochs. For working memory encoding, we examined beta power during the 500-1000ms after the word first appeared on the screen. For feedback we examined the 250-750ms after feedback first appeared. We averaged beta power within these time periods for correct and incorrect trials on an individual channel basis. We also computed resting beta power on an individual channel basis and averaged it over the full two-minute recording period.

Statistical analysis was performed in MATLAB. To examine changes in power with working memory encoding and feedback we used the Wilcoxon signed-rank test to compare beta power during working memory encoding and feedback to beta power during the 500ms baseline period just prior to stimulus and feedback onset. We also used the Wilcoxon signed-rank test to assess the mean beta power differences between correct and incorrect trials during encoding and feedback, to determine whether these changes were related to task performance. To examine the relationship between beta power changes and cognitive function, we compared caudate and DLPFC beta power averaged during working memory encoding and feedback for correct trials on an individual channel basis, as well as resting-state beta power, between cognitively impaired and non-impaired patients using Wilcoxon signed-rank tests. We used Pearson correlation to evaluate the relationship between each subject’s composite cognitive domain scores and beta power during working memory encoding, feedback, and at rest averaged on an individual subject basis. We also used Pearson correlation to examine the relationship between each subject’s caudate volume normalized by intracranial volume and their task-based and resting-state beta power.

Lastly, we conducted a multivariate analysis using each subject’s UPDRS Off score, Levodopa equivalent dose, caudate volume, and caudate or DLPFC beta power during resting-state, working memory encoding, and working memory feedback as predictor variables for preoperative memory function and task performance using normally-distributed generalized linear regression models.

## Results

Fifteen subjects participated in the study. Thirteen subjects were male and two were female, fourteen were right-handed and one was left-handed, ten underwent bilateral STN implants and five underwent bilateral GPi implants. Mean age at time of surgery was 63±6.6 years (mean ± standard deviation), mean disease duration was 8.4±2.9 years, mean preoperative UPDRS off medication score was 45.8±12.0 while mean preoperative UPDRS on medication score was 20.4±12.1, and mean preoperative Levodopa equivalent dose was 1407.9±601.2. Fourteen subjects completed both resting-state data collection and the verbal working memory task, while one subject completed only resting-state data collection. Across all subjects, 25 caudate and 50 DLPFC channels were included in the resting-state analysis, while 21 caudate and 50 DLPFC channels were included in our task-based analysis. For the fourteen patients that completed the 2-back task, mean task performance was 75.8±11.7% correct, 21.2±8.7% incorrect, 3.0±4.3% unanswered. These findings are summarized in **Table 1** and **Figure 2**.

**Table 1.** Patient Demographic and Disease Related Information.

| Patient Number | Age (years) | Sex | Race | Handedness | Duration of Disease (years) | Target | Caudate Channels | Cortex Channels | UPDRS score - off | UPDRS score - on | Levodopa equivalent dose |
| --- | --- | --- | --- | --- | --- | --- | --- | --- | --- | --- | --- |
| 1 | 56 | M | White | Right | 5 | STN, bilateral | 2 | 3 | 49 | 30 | 1400 |
| 2 | 68 | M | White | Right | 9 | STN, bilateral | 4 | 0 | 35 | 9 | 800 |
| 3 | 62 | M | White | Right | 9 | STN, bilateral | 3 | 2 | 53 | 28 | 1240 |
| 4 | 60 | F | White | Right | 11 | STN, bilateral | 3 | 3 | 49 | 35 | 1600 |
| 5 | 58 | M | White | Right | 11 | STN, bilateral | 0 | 6 | 60 | 5 | 2450 |
| 6 | 62 | M | White | Left | 5 | STN, bilateral | 2 | 3 | 41 | 10 | 2050 |
| 7 | 56 | M | White | Right | 6 | STN, bilateral | 2 | 3 | 71 | N/A | 100 |
| 8 | 64 | M | White | Right | 8 | STN, bilateral | 2 | 3 | 30 | 2 | 1900 |
| 9 | 60 | M | White | Right | 7 | STN, bilateral | 3 | 0 | 32 | 24 | 1200 |
| 10 | 66 | M | White | Right | 9 | GPI, bilateral | 0 | 6 | 44 | 28 | 500 |
| 11 | 56 | M | White | Right | 15 | GPI, bilateral | 0 | 5 | 47 | 25 | 1300 |
| 12 | 57 | M | White | Right | 5 | GPI, bilateral | 0 | 5 | 33 | 12 | 1945 |
| 13 | 76 | M | White | Right | 9 | GPI, bilateral | 2 | 3 | 60 | 45 | 2000 |
| 14 | 67 | M | White | Right | 5 | STN, bilateral | 2 | 3 | 30 | 10 | 1333.34 |
| 15 | 77 | F | White | Right | 12 | GPI, bilateral | 0 | 5 | 53 | 23 | 1300 |
| Total/Avg | 63.0 | 13M/2F | White | 14R/1L | 8.4 | 10STN/5GPI | 25.0 | 50.0 | 45.8 | 20.4 | 1407.9 |

**Figure 2.**
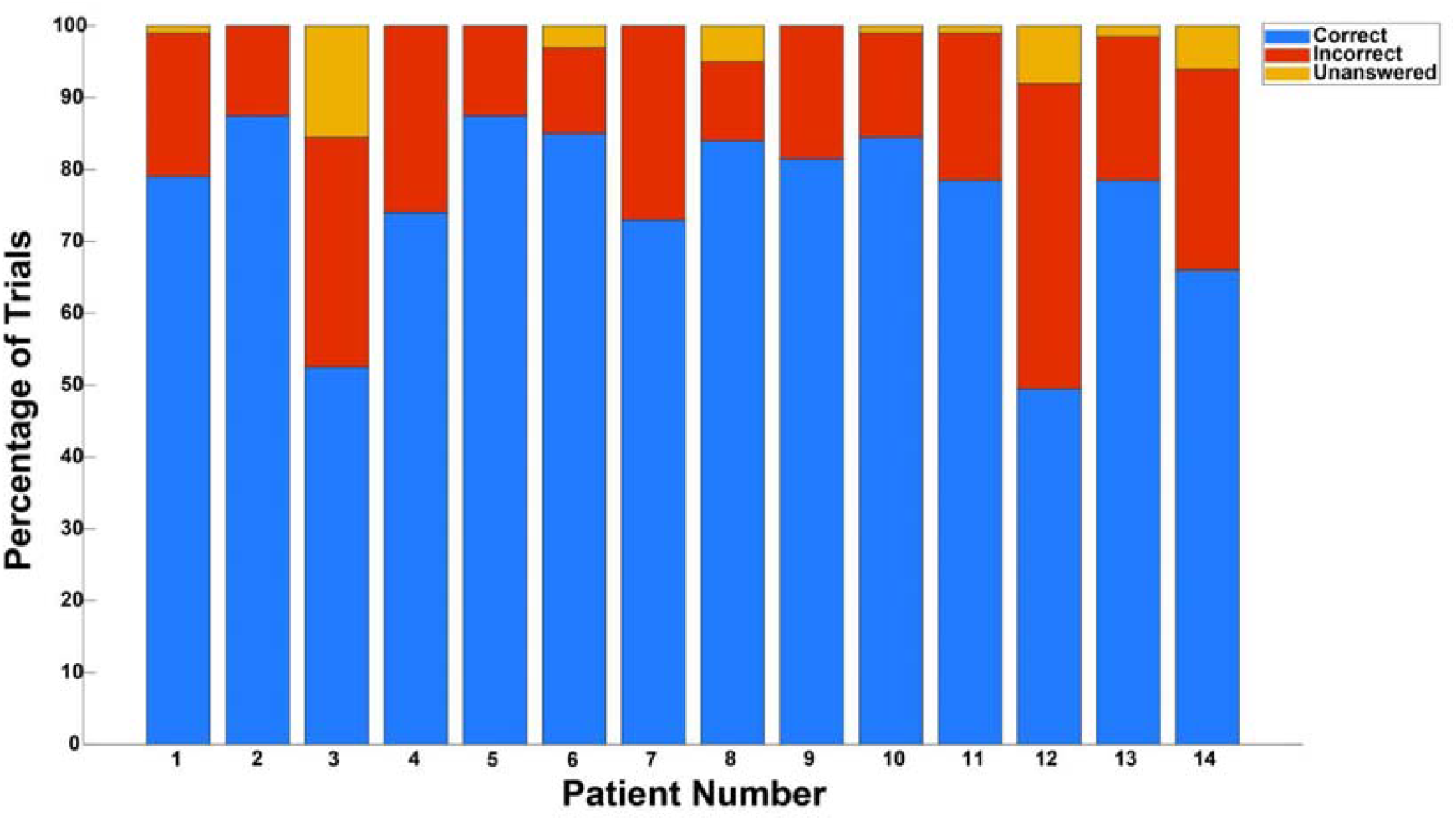
Task Performance By Subject. Percentage of correct (blue), incorrect (red), and unanswered (yellow) trials per subject, averaged across blocks.

### Working Memory Related Beta Power Changes

Both caudate and DLPFC exhibited a decrease in beta power during working memory encoding, with average beta power during this period for words subsequently recalled correctly significantly lower than during baseline [p=0.0026 and p=0.0039, respectively] (**Figure 3**). There was also a significant difference between average beta power for correct and incorrect trials for both caudate and DLPFC [p=0.0051 and p=5.2E-04, respectively] (**Figure 5)**. In caudate, there was greater beta power decrease for correct than incorrect trials, whereas the opposite was true for DLPFC (**Figure 5**).

**Figure 3.**
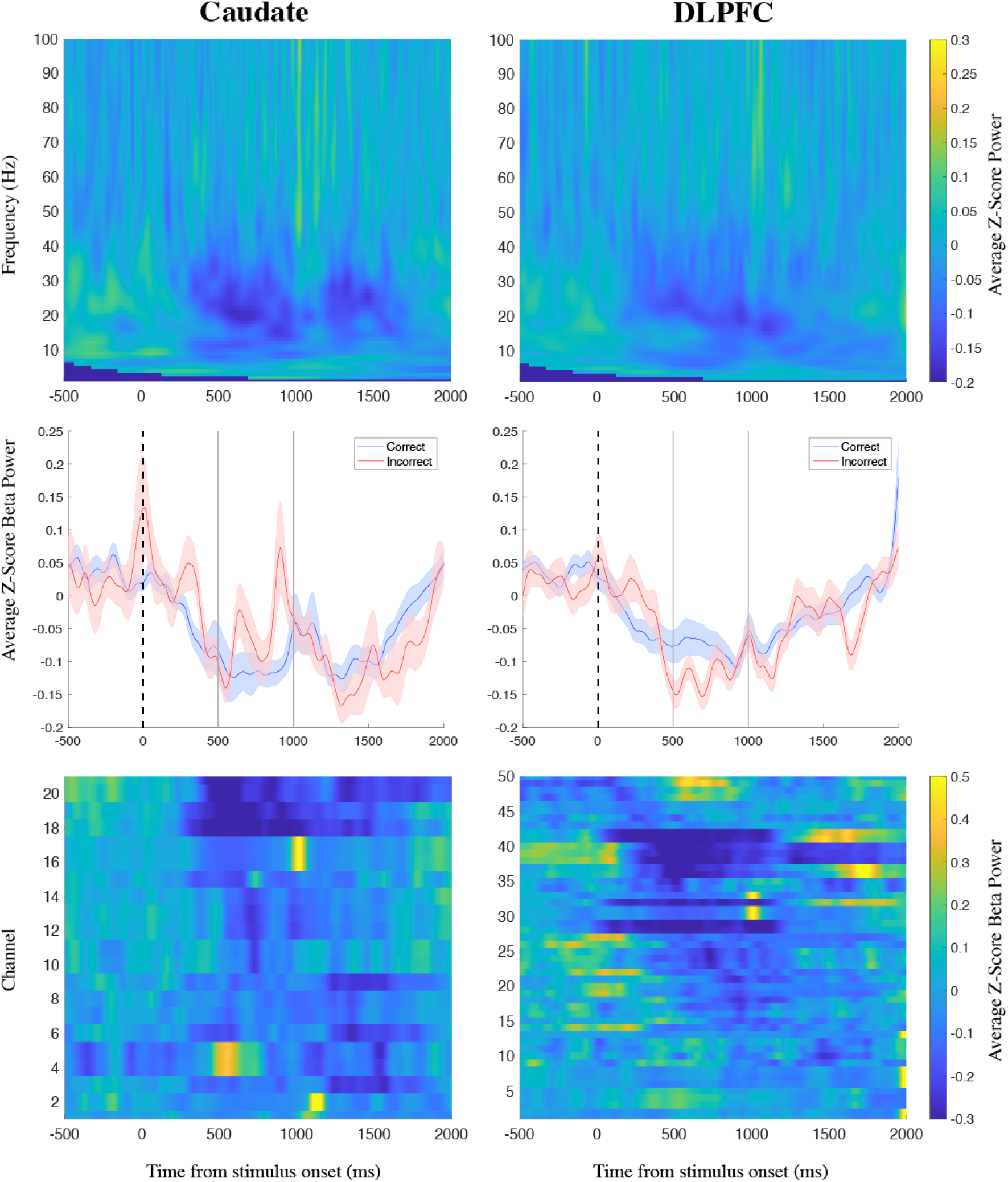
Caudate and DLPFC Power During Encoding. (Top) Spectrograms of z-scored caudate and DLPFC power averaged over correct trials, aligned to stimulus onset. (Middle) Time courses of caudate and DLPFC beta power averaged across subjects on a channel basis during correct and incorrect trials. (Bottom) Changes in z-scored beta power across channels during working memory encoding of correct trials. For correct trials, there was a significant decrease in power during working memory encoding 500-1000 ms following stimulus onset centered in the beta band for both caudate and DLPFC.

Average caudate and DLPFC beta power increased significantly compared to baseline during the 250 to 750ms following feedback presentation for trials where subjects responded correctly [p=0.0010 and p=0.0013] (**Figure 4**). There was no significant change in caudate or DLPFC beta power following feedback for incorrect trials. There was a significant difference between average beta power for correct and incorrect trials in both caudate and DLPFC during this period [p=1.1E-04 and p=4.2E-05] (**Figure 5)**. For both caudate and DLPFC, average beta power exhibited greater increases after correct trials than after incorrect trials (**Figure 5)**.

**Figure 4.**
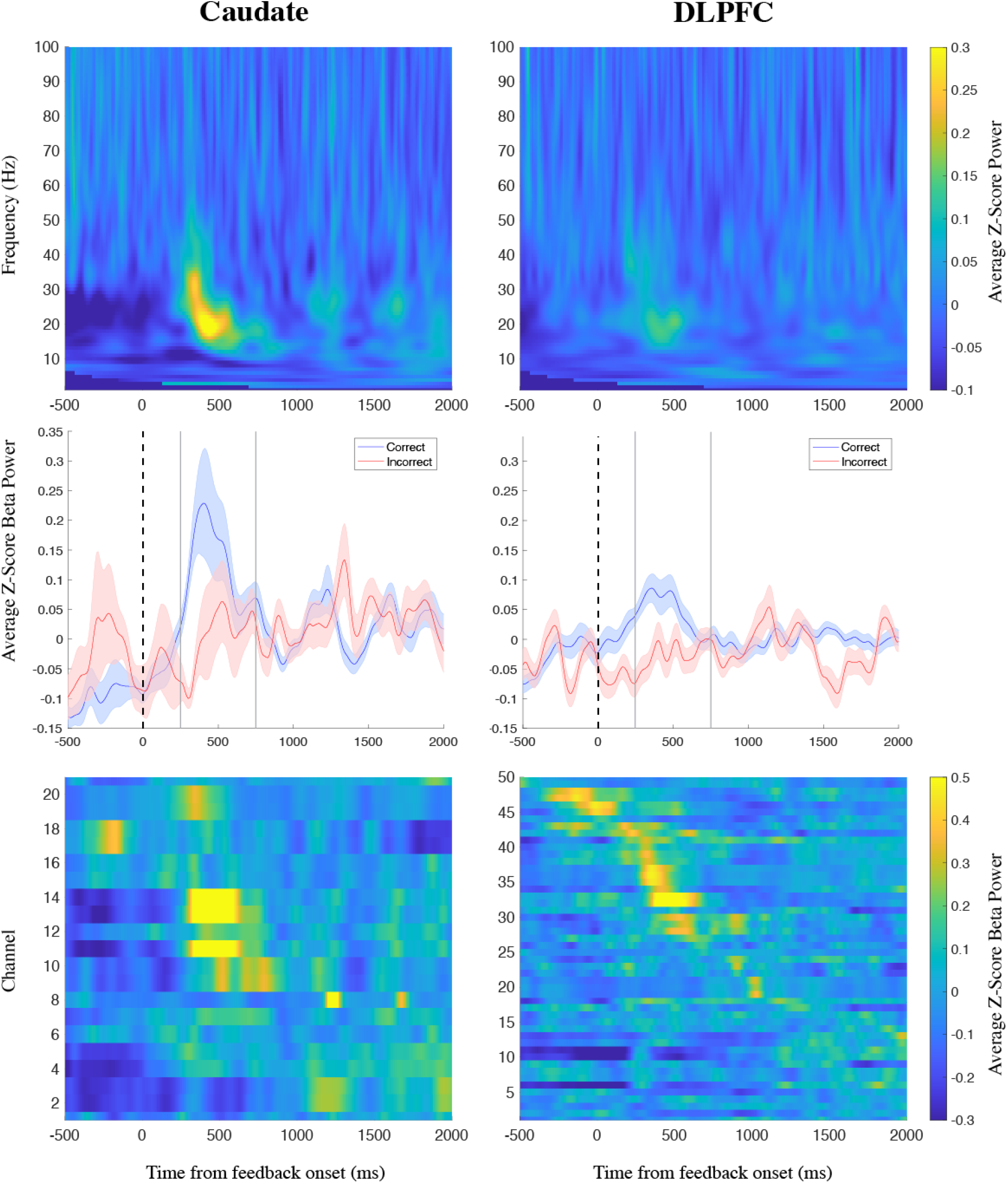
Caudate and DLPFC Power During Feedback. (Top) Spectrograms of z-scored caudate and DLPFC power averaged over correct trials, aligned to feedback onset. (Middle) Time courses of caudate and DLPFC beta power averaged across subjects on a channel basis during correct and incorrect trials. (Bottom) Changes in z-scored beta power across channels during feedback for correct trials. There was a significant increase in power during 250-750 ms following feedback for correct trials, centered in the beta band for both caudate and DLPFC.

**Figure 5.**
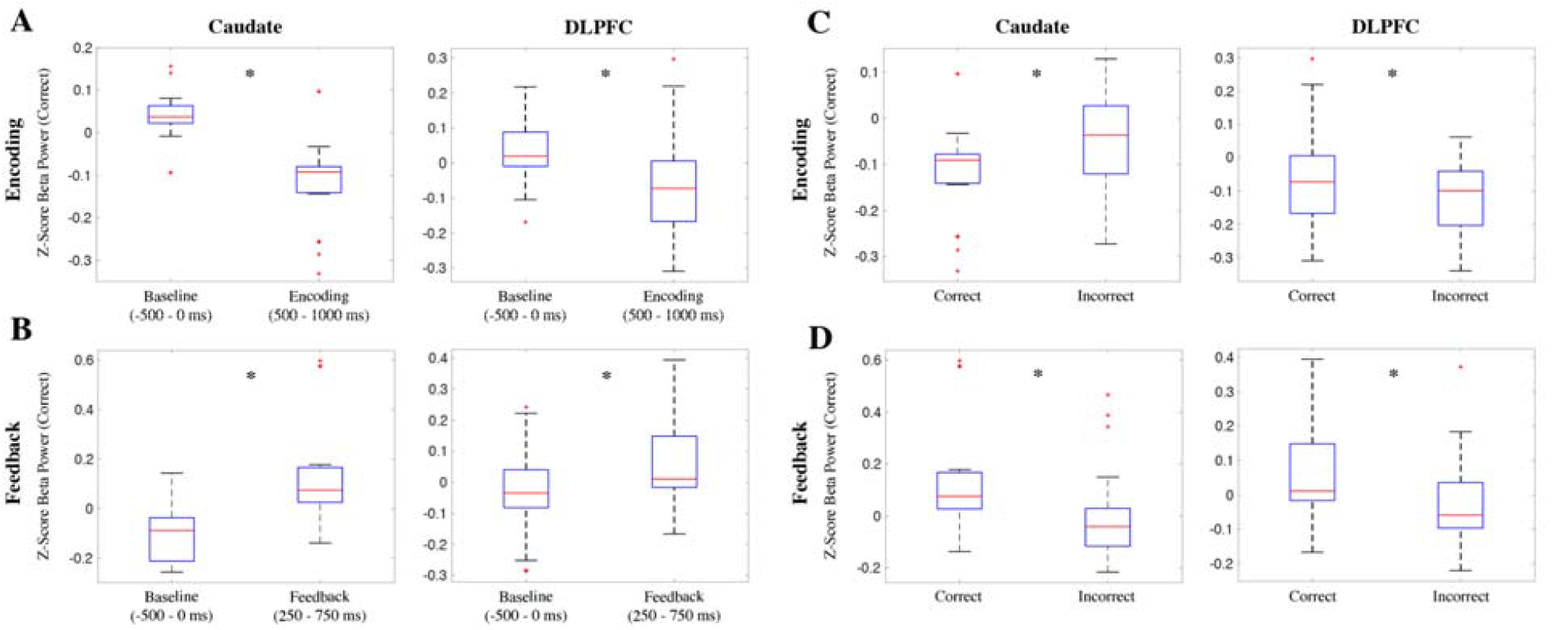
Caudate and DLPFC Beta Power Differences During Task. (A) Average caudate and DLPFC beta power during baseline compared to during working memory encoding. Both caudate and and DLPFC beta power significantly decreased during encoding. (B) Average caudate and DLPFC beta during baseline compared to during feedback. Both caudate and and DLPFC beta power significantly increased during feedback. (C) Average caudate and DLPFC beta power during encoding for correct and incorrect trials. Caudate DLPFC beta power was significantly lower for correct trials, while DLPFC beta power was significantly lower for incorrect trials. (D) Average caudate and DLPFC beta power during feedback for correct and incorrect trials. Both caudate and DLPFC beta power were significantly greater during correct trials. Asterisks denote significance.

There was no correlation between average caudate and DLPFC resting-state beta power and beta power during either encoding or feedback for correct trials. Likewise, neither reaction time nor caudate volume showed a significant correlation with caudate and DLPFC beta power during resting-state, correct trial encoding, or correct trial feedback.

### Relationship between beta power and cognitive impairment

Caudate and DLPFC beta power decreases during working memory encoding were related to cognitive function, with subjects with cognitive impairment having smaller beta power decreases during encoding compared to those with normal cognition [p=1.9E-04 and p=2.5E-03] (**Figure 6)**. We found a significant negative correlation between average caudate beta power during encoding for correct trials and preoperative composite memory domain cognitive scores [R=-0.77, p=0.026], with a greater encoding-related decrease in beta power corresponding with better memory function (**Figure 7)**.

**Figure 6.**
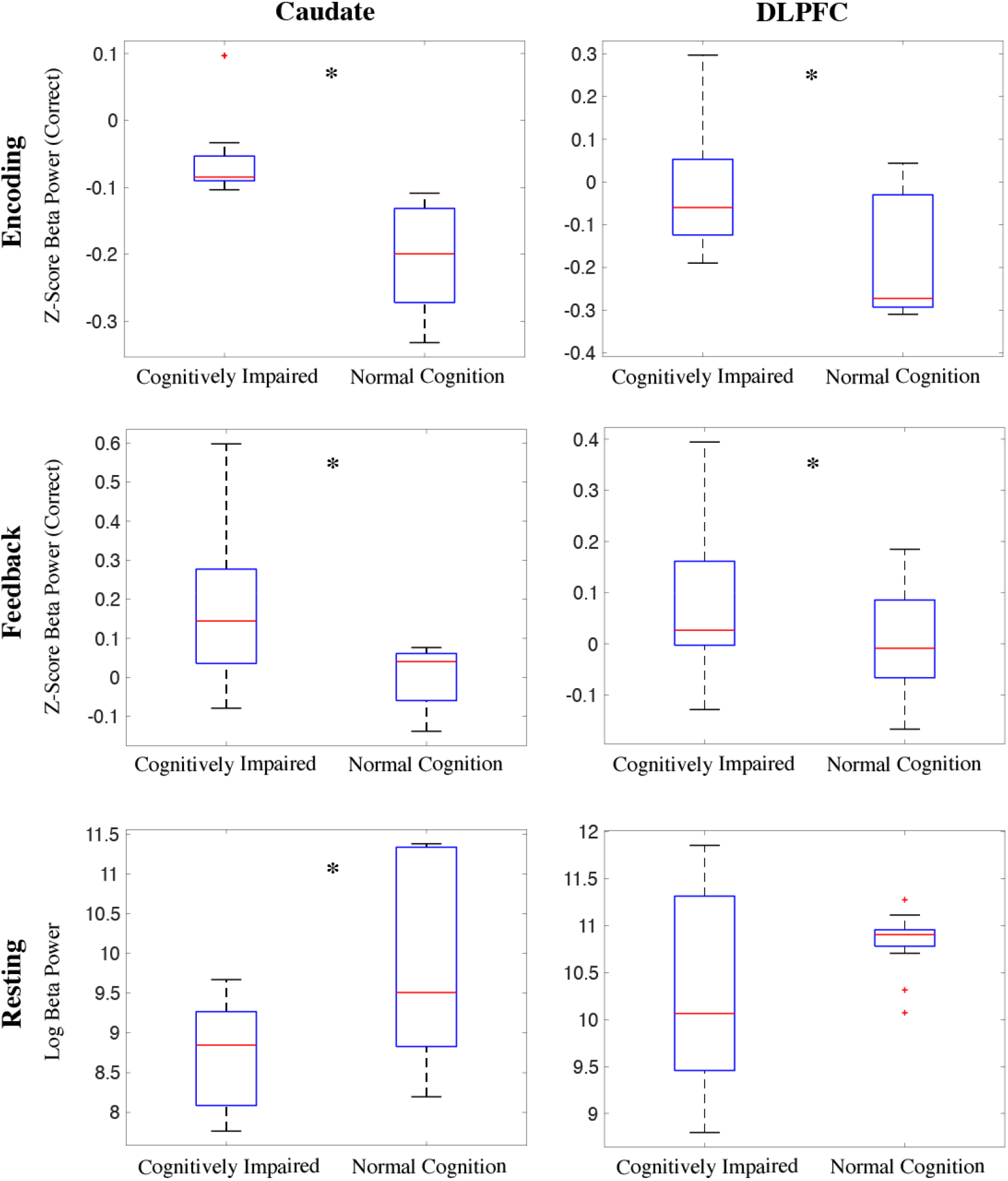
Caudate and DLPFC Beta Power Differences Between Subjects With Cognitive Impairment and Subjects With Normal Cognition. (Top) Average caudate and DLPFC beta power during encoding is significantly higher for subjects with cognitive impairment. (Middle) Average caudate and DLPFC beta power during feedback is also significantly higher for subjects with cognitive impairment. (Bottom) Average resting-state caudate and DLPFC beta power is significantly lower for subjects with cognitive impairment. Asterisks denote significance.

**Figure 7.**
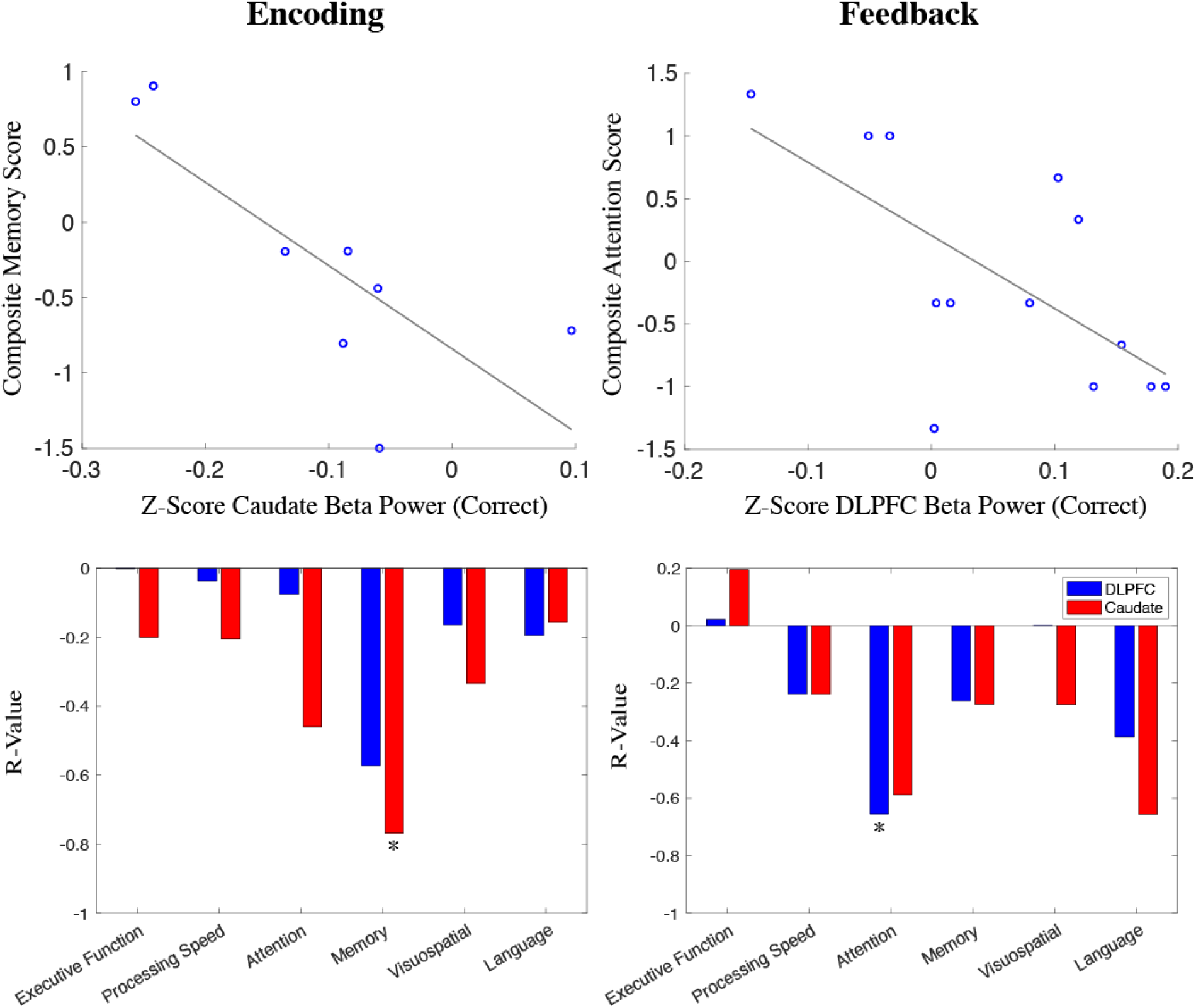
Correlations of Caudate Beta Power and Cognitive Function. (Top) Significant correlations between task-based beta power and domain-specific cognitive function. Caudate beta power during encoding showed a significant negative correlation with preoperative memory function, with greater beta power decreases correlating with better composite memory scores. DLPFC beta power during feedback showed a significant negative correlation with preoperative attention function, with higher beta power increases correlating with worse composite attention scores. (Bottom) R-values of correlations between caudate (red) and DLPFC (blue) beta power during encoding and feedback across all assessed cognitive domains. Asterisks denote significance.

Caudate and DLPFC beta power during feedback for correct trials was also related to cognitive function, with patients with cognitive impairment having greater beta power increases [p=0.027 and p=0.039] (**Figure 6)**. DLPFC beta power during feedback was negatively correlated with preoperative composite attention domain score [R= -0.66, p=0.015], indicating that a greater increase in beta power following task feedback was associated with worse attention (**Figure 7)**.

We found that average resting-state caudate beta power was significantly greater for subjects with normal cognition compared to cognitively impaired subjects [p=0.036], although this difference was not seen for DLPFC beta power. Average caudate and DLPFC resting-state beta power did not exhibit correlation with cognitive function in any domain.

### Multivariate Analysis

A generalized linear regression model using caudate beta power during encoding for correct trials, UPDRS Off score, Levodopa medication dosage, and caudate volume as predictor variables for preoperative memory function [F=4.46, p=0.0417] showed that average caudate beta power during encoding has a significant relationship with subjects’ composite memory scores [p=0.043]. UPDRS Off score, Levodopa medication dosage, and caudate volume were not significant predictors of preoperative memory function.

In contrast, generalized linear regression models using resting-state or caudate feedback beta power, or task-based or resting-state DLPFC beta power, in place of caudate encoding beta power as a predictor variable for preoperative memory function showed no significant relationships with subjects’ composite memory scores. No predictor variables showed a significant interaction with task performance as the response variable.

## Discussion

Analysis of task event-related caudate and DLPFC beta power changes showed that beta power in both structures decreased during working memory encoding for correct trials and increased during task feedback for correct trials. We found that subjects with CI showed a smaller decrease in caudate and DLPFC beta power during encoding and a greater increase in caudate and DLPFC beta power during feedback. Within specific cognitive domains, we found that greater beta power suppression in the caudate during encoding correlated with better preoperative memory function, while higher DLPFC beta power following feedback correlated with greater impairment in the attention domain. resting-state caudate beta power was also significantly lower for subjects with CI. However, there was no correlation between resting-state beta power and domain-specific cognitive impairment or task performance in either the caudate or DLPFC. This is the first study to directly evaluate neurophysiologic human correlates of cognitive CSTC circuits in working memory.

Elevated cortical and basal ganglia beta synchrony has been shown to be a biomarker of PD^19,38^, with research on both human subjects and animal models indicating that dopamine depletion leads to pathologic increases in resting-state beta power.^39–42^ Beta power has been proposed to indicate the active maintenance of a “status quo” state of neuronal activity^38,43,44^, and decreased beta power has been shown to play a crucial role during movement planning and execution.^45^ While the link between elevated beta power in PD and motor symptoms of rigidity and bradykinesia has been well-documented, beta power has also been shown to be involved in cognitive functions such as executive control, working memory, and attention.^43^ Widespread abnormalities in beta band oscillations have also been shown to characterize PD CI, with previous research indicating that global resting-state beta and gamma (30-100 Hz) power is higher in subjects with PD MCI but decreases from PD MCI to Parkinson’s Disease Dementia (PDD).^46^ Lower beta power has also previously been proposed as a strong predictive neurophysiologic marker for PDD.^47^

Our findings confirm previous research on the relationship between cortical beta neural oscillations and working memory encoding. Primate studies demonstrated that lateral PFC beta bursting decreased during working memory encoding and increased following a task response^28–30^, suggesting that beta suppression is necessary for information encoding and that elevated beta activity correlates with a working memory content being cleared out or reallocated once stored information is no longer needed.^43^ Compared to healthy controls, PD subjects have been shown to exhibit both impaired memory strength and reduced global beta power decrease during deep-semantic encoding of a verbal working memory task.^48^ Within the PD subject group, greater beta power suppression in left frontal regions correlated with greater memory strength.^48^ Although there is evidence that event-related beta suppression is involved in the planning and execution of motor tasks^49^, we found no relationship between reaction time and either caudate or DLPFC beta power during encoding. Thus, the beta power suppression we observed during encoding is likely a feature of cognitive function instead of motor action.

While most previous research on the role of beta neural oscillations during working memory focuses on cortical beta activity, our results show that caudate beta power suppression during encoding of correct trials exhibits a stronger correlation with cognitive impairment in the memory domain than DLPFC beta power. This supports that PD cognitive impairment results from the dysfunction of cognitive CSTC circuitry and provides evidence that the striatum gates information to the DLPFC by modulating disinhibition of excitatory CSTC loops.^50^ Computational models have demonstrated that beta power in the basal ganglia can be generated by inhibitory striatal input to the globus pallidus externus (GPe) or excitatory cortical and subcortical inputs to the STN.^43,51^ Drug-naive PD patients exhibit transient underactivation of caudate nuclei, putamen, and globus pallidus during working memory updating.^18^ Although no changes in activation were observed in the DLPFC, basal ganglia under-recruitment disrupts striatal D2-dependent gating mechanisms to cortical regions and contributes to executive and cognitive dysfunction.^18^

Our observations of increased caudate and DLPFC beta power following task feedback are also consistent with previous research into the role of caudate and DLPFC beta power during reinforcement learning.^31^ Both caudate and DLPFC beta power have been shown to increase significantly following feedback for correct trials during a learning task, with DLPFC beta power correlating with learning over time.^31^ There is evidence that caudate dopamine binding correlates with learning-related activation in the DLPFC, which then allows for DLPFC to optimize task performance.^52^ However, the correlation we found between attention impairment and DLPFC beta power following correct trial feedback suggests that caudate beta power in this case may not be directly attributable to successful reinforcement learning.

Our finding that resting-state caudate beta power was significantly lower for subjects with cognitive impairment supports previous research on decreased beta power serving as a predictor for severe cognitive impairment including PDD, although it is not consistent with findings that resting-state beta power is elevated in PD MCI before decreasing with the onset of PDD.^46^ These results further support the role of CSTC circuitry in cognition, with dysfunction in corticostriatal connections potentially underlying lower resting-state caudate beta power in PD CI.

Although previous research has suggested that lower caudate volume is linked with greater beta power, we did not find any significant relationship between caudate volume either task-based or resting caudate and DLPFC beta power.^53^

Understanding of the neurophysiologic biomarkers and dynamics associated with PD CI can clinically translate into development of neuromodulation interventions for patients with medically refractory PD and comorbid CI to improve quality of life and reduce disease-related mortality. Newer technological advancements in DBS devices allow for closed-loop adaptive or responsive stimulation, applying stimulation only at times when a certain designated biomarker is present, which prolongs battery life and minimizes stimulation-related side effects. Several device systems have closed-loop stimulation capabilities that leverage beta power peaks within GPi or STN, noted to correlate with PD motor symptom severity, to initiate timing and cessation of stimulation.^54^ Analogously, this closed-loop capability could be applied to cognitive CSTC circuit structures such as the caudate or DLPFC with electrodes that span both motor and cognitive targets. Other studies have reported efficacy of DBS for memory enhancement, although with alternative targets such as mesial temporal lobe structures^55^ and the hypothalamus/fornix.^56^ However, the efficacy of targeting cognitive CSTC structures for memory improvement in CI patients remains to be determined.

There are several limitations of our study worth mentioning. First, our sample size is small with 15 patients and 75 channels total, which limits the power of our study to detect differences between subgroups. Additionally, channels with significant artifact or channels out of the target region on post-operative imaging were excluded, which leads to inevitable selection bias. There is no control group comparison to healthy subjects, which limits our ability to generalize these results outside of PD patients or to determine features of PD-specific CI. We also limited our analysis to the beta band given prior research demonstrating its value in PD motor symptoms, learning, and memory. LFPs have inherent limitations in spatial resolution of neural signaling and could be improved with single unit recordings. Furthermore, patients with severe cognitive impairment are generally considered to be poor surgical candidates for DBS due to the perioperative risk of potentially worsening CI and because of the need for patients to comply with device self-management and clinical follow up, which limits our ability to capture the entire clinical spectrum of PD CI. Lastly, patients are off of all PD medications for surgery, which confounds their neural signals as well as our ability to assess their “native” cognitive state while on medications.

Our findings demonstrate that caudate and DLPFC beta power changes occur during working memory and correlate with cognitive function in PD patients. These findings support a role for cognitive CSTC beta power changes in PD CI. Future studies may examine whether these changes are dopamine mediated and whether cognitive beta power changes may be a target for neuromodulation for PD cognitive symptoms.

## Conclusion

Dopaminergic depletion in PD leads to altered neurophysiology within the cognitive CSTC circuit structures such as the caudate and DLPFC, which may contribute to working memory deficits and PD CI. Understanding of neurophysiologic correlates of PD CI can inform development of novel neuromodulatory interventions for medically refractory PD patients with comorbid CI.

## Supplementary

### Resting Beta Power of Cognitively Impaired and Normal Cognition Groups

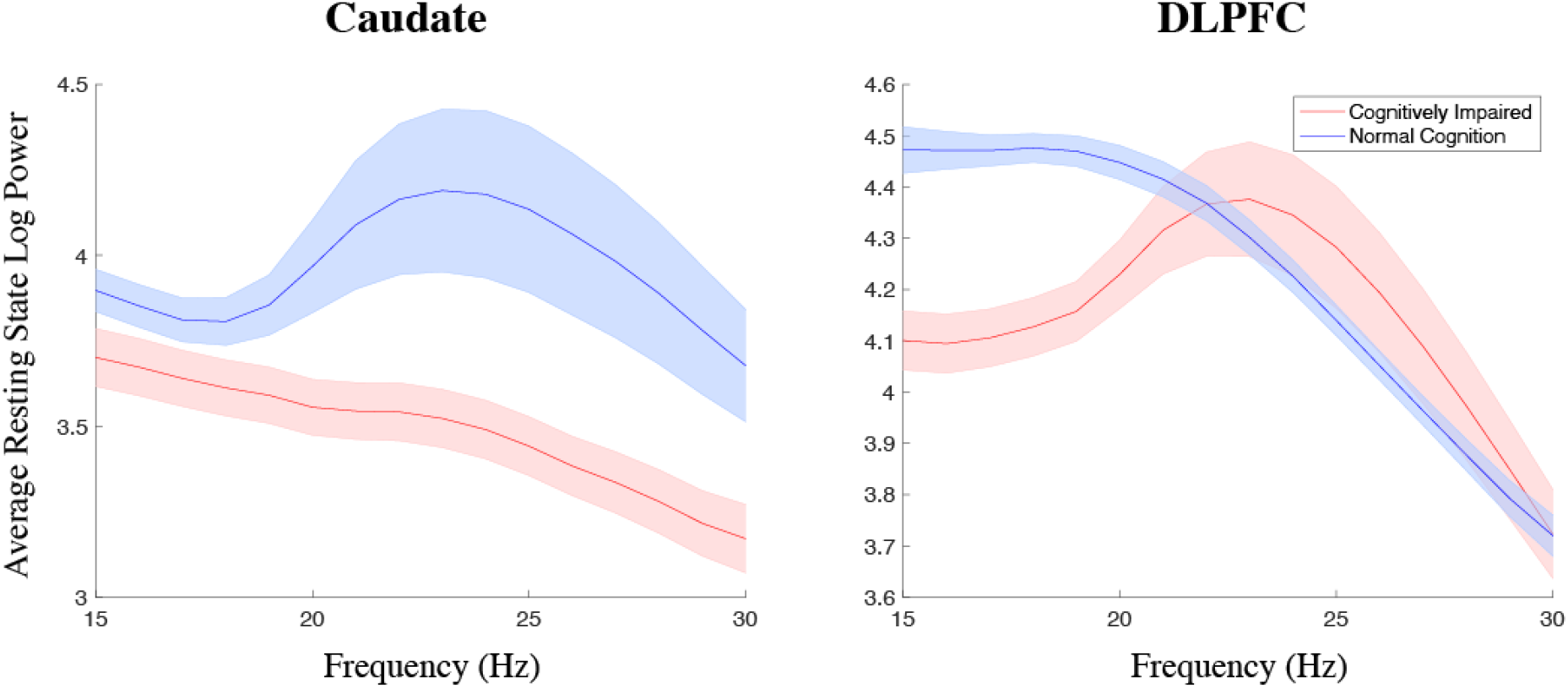

### Demographic Comparisons Between Cognitively Impaired and Normal Cognition Groups

|  | Age (Years) | Disease<br>Duration<br>(Years) | UPDRSOff | UPDRSON | Levodopa<br>Equivalent<br>Dose | Caudate<br>Volume (ICV) | Task<br>Performance (%) |
| --- | --- | --- | --- | --- | --- | --- | --- |
| Wilcoxon Rank Sum P-value | 0.93906094 | 0.43836164 | 0.54505495 | 0.32500833 | 0.47392607 | 0.27212787 | 0.36363636 |
| Cognitively Impaired Mean | 63.7777778 | 7.77777778 | 47.7777778 | 23.125 | 1421.66667 | 0.41354386 | 72.4949264 |
| Cognitively Impaired SD | 8.01214356 | 2.68224616 | 12.813838 | 14.307216 | 695.997486 | 0.03979422 | 13.7980495 |
| Normal Cognition Mean | 61.8333333 | 9.33333333 | 42.8333333 | 16.8333333 | 1387.22333 | 0.39486072 | 81.9095637 |
| Normal Cognition SD | 4.83390801 | 3.44480285 | 12.3193615 | 9.9079093 | 555.259236 | 0.04056238 | 5.43641334 |

### Neuropsychology tests

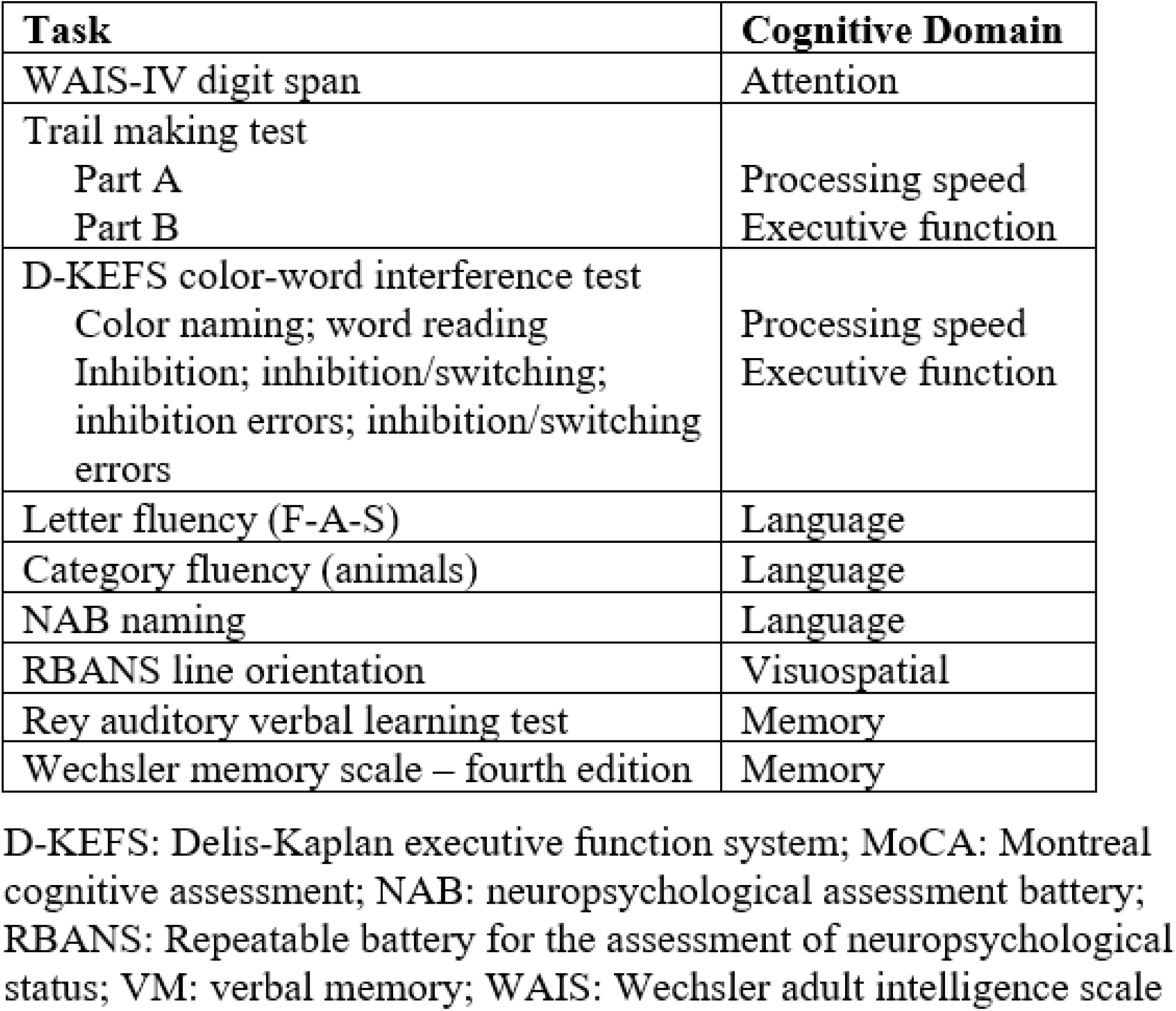

## Notes

### Competing Interest Statement

The authors have declared no competing interest.

### Summary of Updates

Manuscript text, updated analysis, additional figures

